# Human white matter myelination rate slows down at birth

**DOI:** 10.1101/2023.03.02.530800

**Authors:** Mareike Grotheer, David Bloom, John Kruper, Adam Richie-Halford, Stephanie Zika, Vicente A. Aguilera González, Jason D. Yeatman, Kalanit Grill-Spector, Ariel Rokem

## Abstract

The formation of myelin, the fatty sheath that insulates nerve fibers, is critical for healthy brain function. A fundamental open question is what is the impact of being born on myelin growth. To address this question, we evaluated a large (n=300) cross-sectional sample of newborns from the Developing Human Connectome Project (dHCP). First, we developed new software for the automated identification of 20 white matter bundles in individuals that is well-suited for large samples. Next, we fit linear models that quantify T1w/T2w, a myelin-sensitive imaging contrast, increases along bundles. We found faster growth of T1w/T2w along the lengths of all bundles before birth than right after birth. Further, in a separate longitudinal sample of preterm infants (N=34), we found lower T1w/T2w at term-equivalent age than in full-term peers. By applying the linear models fit on the cross-section sample to the longitudinal sample of preterm infants, we find that their delay in T1w/T2w growth is well explained by the amount of time preterm infants spend developing in utero and ex utero. These results suggest that being born slows the rate of myelin growths. This reduction in the rate of myelin growth at birth, in turn, explains lower myelin content in individuals born preterm, and could account for long-term cognitive, neurological, and developmental consequences of preterm birth. We hypothesize that closely matching the environment of infants born preterm to what they would have experienced in the womb may reduce delays in myelin growth and hence improve developmental outcomes.

---

The human brain develops most rapidly early in life – in fact, the total brain volume increases by 101% in the first year of life alone (1). The large-scale structural features of the brain, such as the major gyri and sulci, as well as the location and trajectory of major white matter tracts, are largely established by birth (2). Nevertheless, the dramatic brain volume increases in early life are accompanied by various important and tightly regulated microstructural processes, including myelination of the brain’s white matter (3). During myelination, neuronal axons are wrapped in a fatty sheath, which insulates them and thereby enables rapid and synchronized neural communication across the brain. Myelination is a major contributor to brain plasticity and to learning. Thus, abnormalities in myelin development contribute to a plethora of developmental and cognitive disorders (4–6). However, the spatiotemporal pattern and mechanisms that govern myelination at the very onset of life are still poorly understood.

One fundamental open question is what impact birth itself has on the myelination of the brain’s white matter. After all, birth entails a dramatic change in the environment and experiences of the developing human and is associated with a multitude of biological signals. In addition to the null-hypothesis that birth does not impact myelination, there are two developmental hypotheses about how these changes might affect myelination that can be derived from the animal literature (Supplementary Fig. S1). On the one hand, myelination has been shown to be experience-dependent (7–9), which, in turn, would suggest that the dramatic increase in incoming sensory information at birth, such as the increase in visual input, provokes an increased rate of myelination *ex utero* compared to *in utero*. On the other hand, the protective environment of the womb, with its consistent supply of nutrients, oxygen, warmth, etc. could promote rapid myelination. After all, myelination is tightly coupled with nutrition (10–12) and sensitive to fluctuations in oxygen supply at specific developmental stages (13). While these hypotheses are not mutually exclusive, they have opposing impact on myelin development. Distinguishing between these hypotheses is particularly critical in the context of preterm birth, as preterm birth changes the proportion of time an infant develops *in utero* and *ex utero* and has been shown to induce persistent white matter differences, also associated with long-term cognitive consequences (14). Importantly, the impact of birth on myelin development may also differ between infants born preterm and those born full-term. This is because the less mature preterm brain is thought to be particularly vulnerable and to benefit more from the protective environment of the womb than the full-term brain (15, 16).

One way to distinguish between these developmental hypothesis is to fit models that take into account both the time spent *in utero* and the time spent *ex utero* and to then compare the effect of each of these periods on myelin growth (17). Fitting such models, however, requires collecting large data sets – either sampling the same individuals multiple times, or sampling a large group of individuals, with substantial variance in gestational age at birth and time between birth and measurement. As such, this type of question cannot be fully addressed using *ex vivo* dissection studies (such as (18–21)), which compare myelin development across individuals using labor-intensive and user-dependent methods, and are limited to small samples that are often associated with brain pathologies (22). Previous *in vivo* studies of early life white matter development (3, 23–27) largely focused on diffusion-based metrics (e.g., fractional anisotropy). These metrics, while highly reliable representations of the measured signal (28), are not specific to myelination. Instead, they reflect various changes to the white matter including the diameter, spacing, and orientation of fibers along with myelination (29). More direct *in vivo* estimates of myelination can be obtained using specialized quantitative metrics (30–36) such as myelin water fraction (MWF) or the longitudinal relaxation rate of the MR signal, R1. Unfortunately, acquiring quantitative MWF or R1 data is time-consuming and such studies are not only rare in early life samples (for exceptions see: (37–39)), but also conducted at a smaller scale as the one needed for the current experimental question. Fortunately, measurements of R1 can also be approximated using more commonly obtained MRI sequences. In fact, T1-weighted (T1w) and T2-weighted (T2w) imaging sequences are both sensitive to myelin content, but their sensitivity is inverse. This means that instrumental factors (such as B1+ field gain), which affect both of these measurements equally, can be canceled out by dividing the T1w image by the T2w image (40). Indeed, the T1w/T2w ratio is correlated with R1 in adult cortex (41) and is linked to myelination in infants, as well (42). This is particularly exciting as T1w and T2w data are included in large early-life datasets, enabling us to explore the impact birth has on white matter myelination. To distinguish developmental hypotheses on the impact of birth on white matter myelination, first, we developed and shared open-source software (baby automated fiber quantification in python (pyBabyAFQ)) that enables the identification of 20 white matter bundles in individual infants and the analysis of T1w/T2w and other measures along their lengths on a large scale. We then used this software to analyse newborn data provided by the developing human connectome project (dHCP (43)). We divided this data into two independent samples. The “cross-sectional sample” included 300 infants with large variations in gestational ages at birth (range: 25–42 weeks, delta: 17 weeks) and time between birth and measurement (range: 0 –17 weeks) allowing us to model T1w/T2w growth in the weeks before and after birth and to hence assess the impact of birth on T1w/T2w increase. We also used the models fit on the cross-sectional sample to predict T1w/T2w in a “longitudinal sample” that contained 34 preterm infants scanned once shortly after birth and once at >40 weeks post-menstrual age, as well as a matched group of full-term infants. This allowed us to directly relate the observed impacts of birth to T1w/T2w measured in preterm infants.

Our analyses showed faster rate of development of T1w/T2w along the lengths of white matter bundles *in utero* compared to *ex utero*, suggesting that myelination rate slows down right after birth. We also provide evidence that delays in T1w/T2w development observed in infants that are born preterm are explained by the differential T1w/T2w growth rates in utero and ex utero and the infants’ shorter duration developing *in utero*. Our work hence offers a mechanistic explanation for preterm birth related developmental delays based on the early occurrence of birth, which may suggest that providing a “womb-like” environment could reduce preterm birth related delays in myelin development.

## Results

### T1w/T2w varies with post-menstrual age at scan during early infancy

To evaluate the relative contributions of time spent *in utero* and *ex utero* on the myelination of the human infant’s white matter, first we developed a fully-automated, easy to use, open-source, and scalable software package for the identification of white matter bundles in individual infant’s brains. This software, called “baby automated fiber quantification in python (pyBabyAFQ)” is specifically designed for the infant brain and identifies 20 white matter bundles in each individual’s native brain space (see Fig. 1 and Supplementary Fig. S2 for example bundles). PyBabyAFQ was validated against manually identified bundles and shown to outperform commonly used software developed for adult data (see SI results and Supplementary Fig. S3). Using pyBabyAFQ, we evaluated changes in T1w/T2w along the length of 20 white matter bundles in individual infants’ brains. We chose T1w/T2w, as we found it to correlate highly with quantitative measures related to myelin (see SI results and Supplementary Fig. S4). We studied two separate samples from the dHCP project: (i) the “cross-sectional sample”, which contained 300 infants scanned shortly after birth (gestational age at birth: range: 24.57 - 42.29 weeks, mean ± SD: 38.30 ± 3.50 weeks; time between birth and scan: range: 0-16.71 weeks, mean ± SD: 1.60 ± 2.49 weeks), and (ii) the “longitudinal sample”, which contained a distinct sample of 34 preterm infants (gestational age at birth: mean ± SD: 31.97 ± 2.96 weeks) scanned once shortly after birth (post-menstrual age at first scan: mean ± SD: 34.34 ± 1.84 weeks) and once at >40 weeks post-menstrual age (age at second scan: mean ± SD: 40.92 ± 1.34 weeks), as well as a group of full-term infants with matched scan ages and genders (post-menstrual age at scan: mean ± SD: 40.92 ± 1.35 weeks). To establish whether T1w/T2w is impacted by the overall amount of time an infant\ developed, we started with the cross-sectional sample and plotted T1w/T2w along the length of 20 bundles separately for three different scan age groups: group 1: scan age 33 thru < 37 weeks; group 2: scan age 37 thru < 41 weeks; group 3: scan age 41 thru < 45 weeks (Fig-1b). We found that T1w/T2w is lowest in group 1, intermediate in group 2, and highest in group 3, supporting the conclusion that post-menstrual age at scan impacts T1w/T2w. Interestingly, T1w/T2w generally appears higher towards the middle of bundles, suggesting more advanced development of the center compared to the periphery of the bundles (note that the sharp peaks at the first and last nodes likely reflect partial volume effects with the gray matter). Moreover, in several bundles (e.g., the cingulum cingulate, CC) the difference between group 1 and group 2 appears more pronounced than the difference between group 2 and group 3, even though the difference in average post-menstrual age at scan between the groups is the same. To further quantify the effect of post-menstrual age at scan on T1w/T2w, next, we fit linear models (LMs) that related T1w/T2w at each node in each bundle to post-menstrual age at scan in the entire cross-sectional sample. On average, the LMs explained 44% of T1w/T2w variance along the lengths of all bundles (R^2^ between: 0.11-0.76, all p values<0.0001, all slopes>0.02, for mean values per bundle see Supplementary Table 1), suggesting that T1w/T2w is positively related to the infants’ post-menstrual age at scan.

**Fig. 1.**
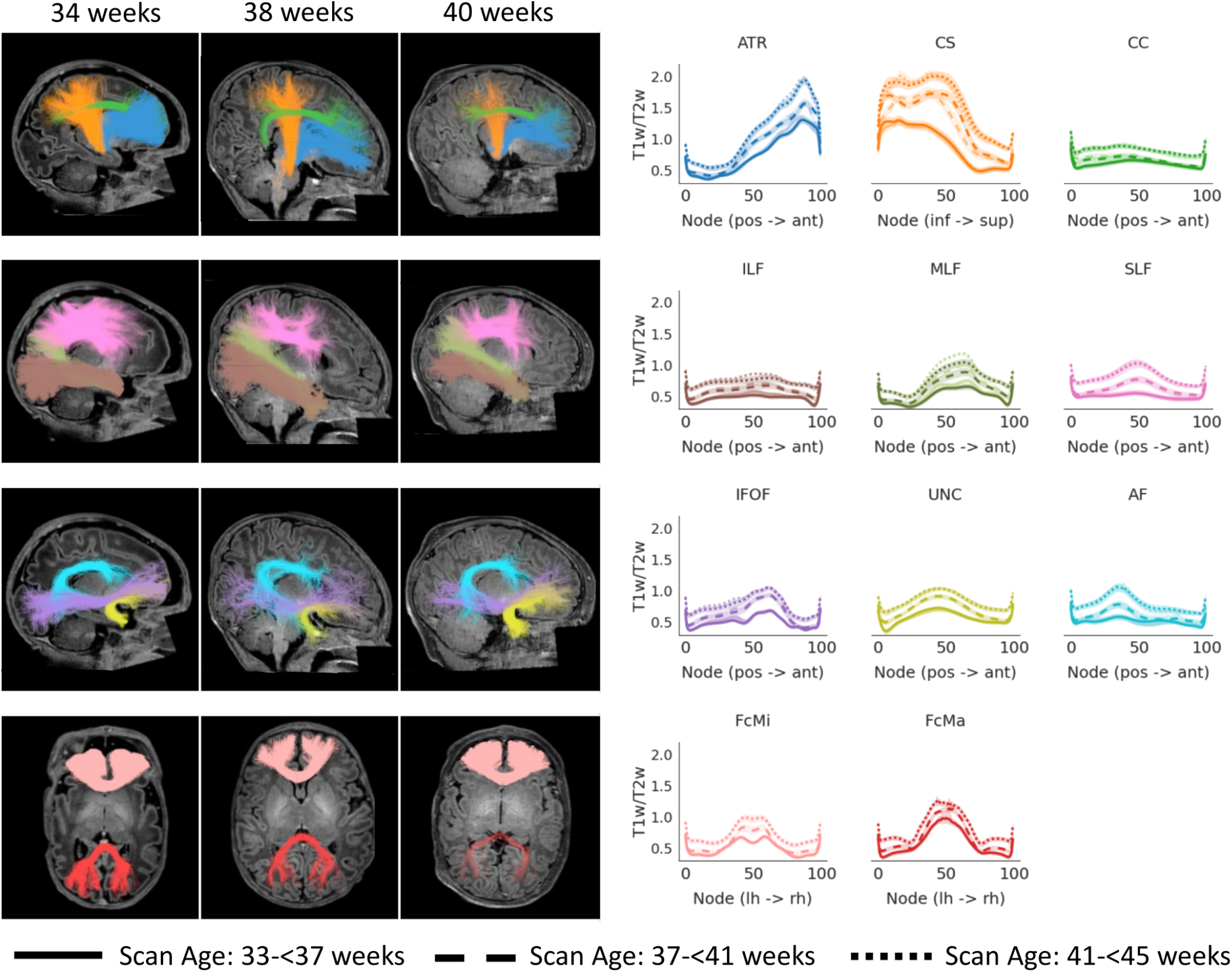
T1w/T2w increases with increasing post-menstrual age at scan. Left: Examples of the evaluated bundles in the native brain space of three infants scanned at 34 weeks, 38 weeks and 42 weeks post-menstrual age, respectively. Right: Line graphs show T1w/T2w along the length of bundles, separately for three scan age groups: group 1: solid line, scan age 33-<37 weeks; group 2: dashed line, 37-<41 weeks; group 3, dotted line, 41-<45 weeks. We find that T1w/T2w increases with increasing post-menstrual age at scan along the lengths of all bundles. For bilateral bundles, both hemispheres are presented in the same subplot, right hemisphere is shown in lighter hue. Shaded regions show 95% bootstrapped confidence intervals. Abbreviations: ATR: anterior thalamic radiation, CS: cortico-spinal tract, CC: cingulum cingulate, FcMi: forceps minor, FcMa: forceps major, ILF: inferior longitudinal fasciculus, MLF: middle longitudinal fasciculus, IFOF: inferior frontal occipital fasciculus, SLF: superior longitudinal fasciculus, UNC: uncinate fasciculus, AF: arcuate fasciculus, pos: posterior, ant: anterior, inf: inferior, sup: superior, lh: left hemisphere, rh: right hemisphere

### T1w/T2w increases more rapidly during in utero development

Importantly, effects of post-menstrual age at scan can be driven by either *in utero* or *ex utero* development. Therefore, we separately tested whether there is a difference in development rate in utero and ex utero (i.e., an effect of being born). In the cross-sectional sample, we fit LMs relating T1w/T2w at each node in each bundle to: (i) gestational age at birth (in utero development), and (ii) time between birth and scan (ex utero development). If being born has no impact on T1w/T2w, then the LMs that separate in utero and ex utero development should fit the data as well as the LMs based solely on post-menstrual age at scan, described above. In contrast, if being born impacts T1w/T2w growth, then the models which separate *in utero* and *ex utero* development should outperform the models based on post-menstrual age alone (due to differences in the slopes for *in utero* and *ex utero* T1w/T2w development). On average, the LMs separating in utero and ex utero development explained 50% of T1w/T2w variance along the lengths of all bundles (R^2^ between: 0.14-0.82, all p values<0.0001, for mean values per bundle see Supplementary Table 1). Model comparison using the Akaike information criterion (AIC) show that the in utero /ex utero models outperform the models based on post-menstrual age at scan in all bundles (AIC difference: mean: 36.03, range: -1-130, mean difference in R^2^: 0.06), suggesting that being born impacts the rate of T1w/T2w change. Further, we found that the rate of T1w/T2w increase is faster *in utero* (mean slope: 0.04; range: 0.02-0.13, for mean values per bundle see Supplementary Table 1) than *ex utero* (mean slope: 0.03; range: 0.002-0.09, for mean values per bundle see Supplementary Table 1), suggesting that, on average, the rate of T1w/T2w development slows down at birth. Indeed, plotting the in utero and ex utero slopes along bundles (Fig. 2) revealed steeper in utero than ex utero rates of T1w/T2w development in all bundles. Interestingly, the magnitude of the observed difference between in utero and ex utero slopes varied between bundles. For example, in projection bundles (e.g. ATR and CS) these differences appear more pronounced than in association bundles (e.g. ILF and MLF) and commissural bundles (FcMi and FcMa). The magnitude of the observed differences in slopes between *ex utero* and *in utero* development even varied along the lengths of a given bundle. For example, at the posterior end of the anterior thalamic radiation (ATR) *in utero* and *ex utero* development rates are more similar than at the anterior end of the bundle. In other bundles, such as the cingulum cingulate (CC), the difference between in utero and ex utero development is comparatively stable along the length of the bundles. The observed difference between in utero and ex utero slopes remained when infants born preterm were excluded from the analysis (Supplementary Fig. S5).

**Fig. 2.**
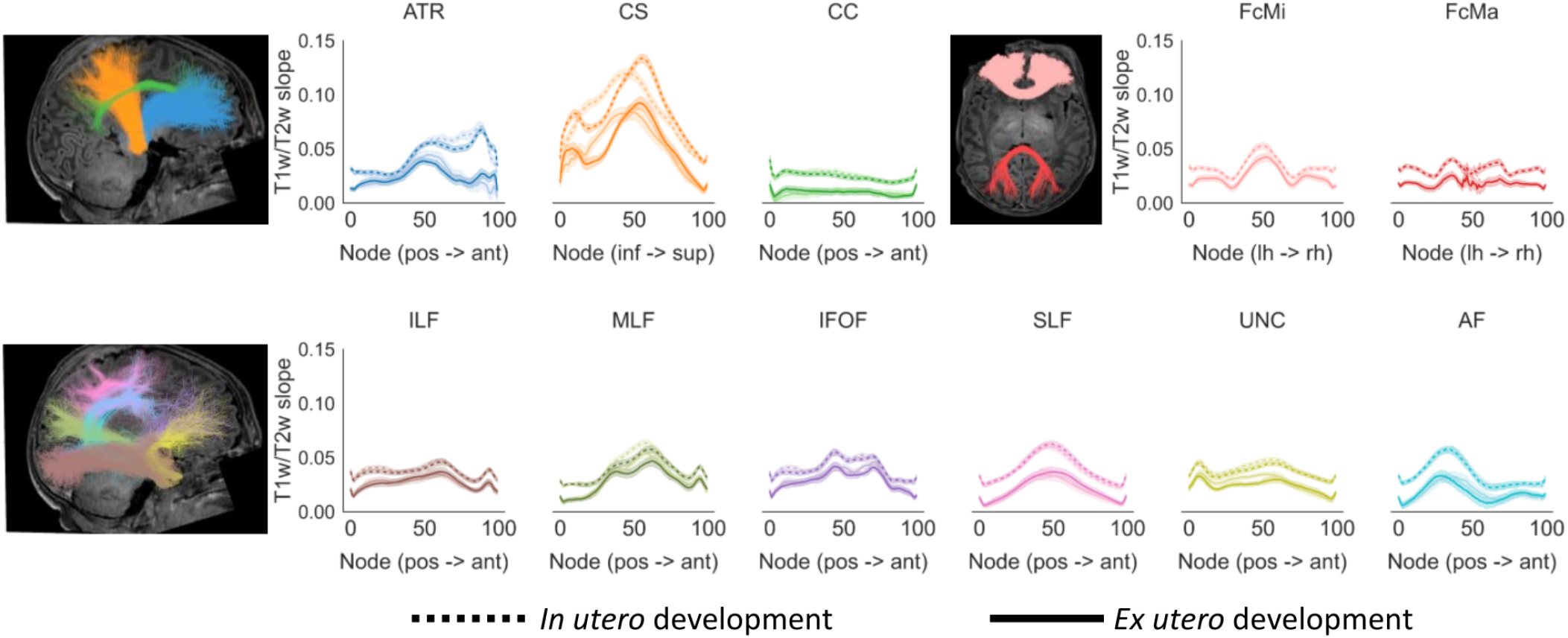
T1w/T2w increases more rapidly in utero compared to ex utero. Brain slices show examples of the evaluated bundles in the native brain space of a representative full-term infant. Line graphs compare the slope (rate of change) of T1w/T2w prior to (in utero development, dotted line) and after (ex utero development, solid line) birth. We find steeper slopes for in utero development along the lengths of all bundles. For bilateral bundles, both hemispheres are presented in the same subplot, right hemisphere is shown in lighter hue. Shaded regions show 95% confidence intervals. Abbreviations: ATR: anterior thalamic radiation, CS: cortico-spinal tract, CC: cingulum cingulate, FcMi: forceps minor, FcMa: forceps major, ILF: inferior longitudinal fasciculus, MLF: middle longitudinal fasciculus, IFOF: inferior frontal occipital fasciculus, SLF: superior longitudinal fasciculus, UNC: uncinate fasciculus, AF: arcuate fasciculus, pos: posterior, ant: anterior, inf: inferior, sup: superior, lh: left hemisphere, rh: right hemisphere

### Infants born preterm show lower T1w/T2w than full-term infants at similar post-menstrual age at scan

Our analyses of a large cross-sectional sample suggest that T1w/T2w development slows down after birth. This, in turn, predicts that infants born preterm will have lower T1w/T2w at term-equivalent age, as they spend less time developing in utero compared to their full-term peers. To test this prediction, next, we focused on the longitudinal sample and compared T1w/T2w between three groups of infants: (i) infants born preterm and scanned shortly after their preterm birth, (ii) the same preterm infants scanned once they reached term-equivalent age, and (iii) a group of full-term infants that were individually matched to each of the preterm infants’ second scan. Matching took into account the infant’s sex and post-menstrual age at scan and after matching, post-menstrual age at scan was 40.92 weeks in both groups. As predicted, preterm infants showed lower T1w/T2w along the lengths of all bundles than their full-term peers when scanned at the same term-equivalent post-menstrual age (Fig. 3). In fact, overall, T1w/T2w of the preterm infants at their second scan is approximately intermediate between T1w/T2w of their first scan and T1w/T2w observed in the full-term group along most bundles, with fine-grained differences between bundles. For example, in the cingulum cingulate (CC) the T1w/T2w measured at the preterm infants’ second scan closely resembled their first scan, suggesting a lack of ex utero growth in T1w/T2w in that bundle. In contrast, in the cortico-spinal tract, the T1w/T2w measured at the preterm infants’ second scan more closely resembled the T1w/T2w measured in the full-term group, suggesting comparatively fast ex utero T1w/T2w growth in that bundle.

**Fig. 3.**
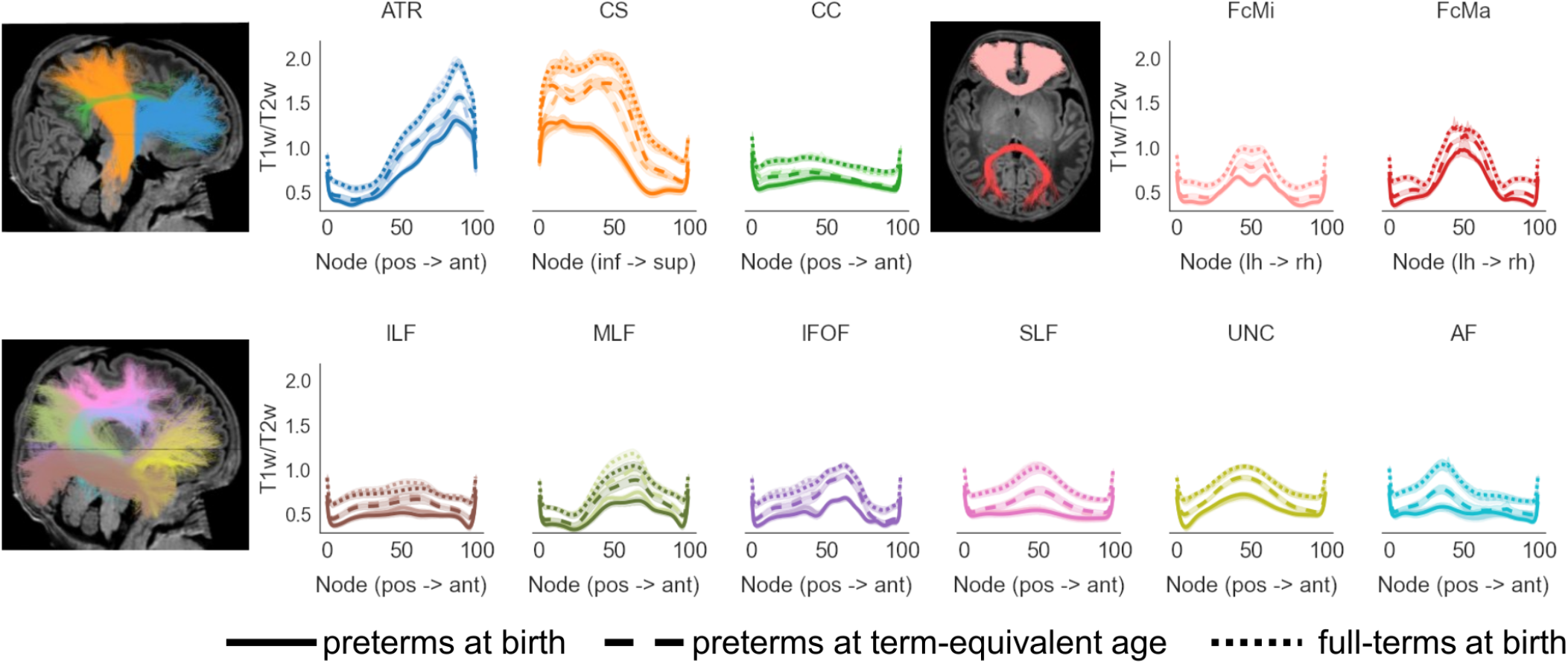
Lower T1w/T2w at term-equivalent age in infants born preterm. Brain slices show examples of the evaluated bundles in the native brain space of a representative preterm infant. Line graphs compare T1w/T2w in three groups: infants born preterm and scanned shortly after their preterm birth (solid line), the same preterm infants scanned once they reached term-equivalent age (dashed line), and a group of full-term infants that was matched 1-on-1 to the preterm infants’ second scan (dotted line). We find lowest T1w/T2w in the preterm infants when scanned shortly after birth, intermediate T1w/T2w at the preterm infants second scan and highest T1w/T2w in the full-term infants. For bilateral bundles, both hemispheres are presented in the same subplot, right hemisphere is shown in lighter hue. Shaded regions show 95% confidence intervals. Abbreviations: ATR: anterior thalamic radiation, CS: cortico-spinal tract, CC: cingulum cingulate, FcMi: forceps minor, FcMa: forceps major, ILF: inferior longitudinal fasciculus, MLF: middle longitudinal fasciculus, IFOF: inferior frontal occipital fasciculus, SLF: superior longitudinal fasciculus, UNC: uncinate fasciculus, AF: arcuate fasciculus, pos: posterior, ant: anterior, inf: inferior, sup: superior, lh: left hemisphere, rh: right hemisphere

### Slower ex utero T1w/T2w development explains lower T1w/T2w in preterm infants

Our analyses thus far suggest that i) development of T1w/T2w slows down at birth, and ii) T1w/T2w is lower in preterm compared to full-term infants at term-equivalent age. In this final analysis, we aimed to explicitly link between these two effects and test if the slower *ex utero* T1w/T2w development explains the lower T1w/T2w in the preterm infants, as they spend more time developing *ex utero*. To this end, we returned to the LMs fit on the cross-sectional sample described above and tested how well these models explain the T1w/T2w measured in the preterm infants at term-equivalent post-menstrual age. Importantly, the models are hence fit and tested on independent samples. The first set of LMs only accounted for post-menstrual age at scan, whereas the second set of LMs accounted for time spend developing *in utero* and *ex utero*. We find that the models based on post-menstrual age at scan overestimate T1w/T2w in the preterm infants at term-equivalent age (Fig. 4a). That is, on average, the predicted T1w/T2w was 0.14±0.04 (±SD) greater than the measured T1w/T2w across bundles. In contrast, the *in utero* /*ex utero* models captured T1w/T2w measured in preterm infants well; on average the predicted T1w/T2w was only 0.03 ± 0.03 (±SD) greater than the measured T1w/T2w across bundles. Accordingly, the models separating *in utero* and *ex utero* development better explained T1w/T2w across all nodes and bundles in each individual preterm infant (post-menstrual age at scan model: mean R^2^: 0.64, range: -0.03 - 0.91, in utero /ex utero model: mean R^2^: 0.87, range: 0.69-0.95; Mann Whitney U-Test comparing R^2^-values: p<0.0001; Fig. 4b). This independent analysis suggests that the lower T1w/T2w observed in preterm infants is largely explained by the fact that they spend more time developing *ex utero* than their full-term peers.

**Fig. 4.**
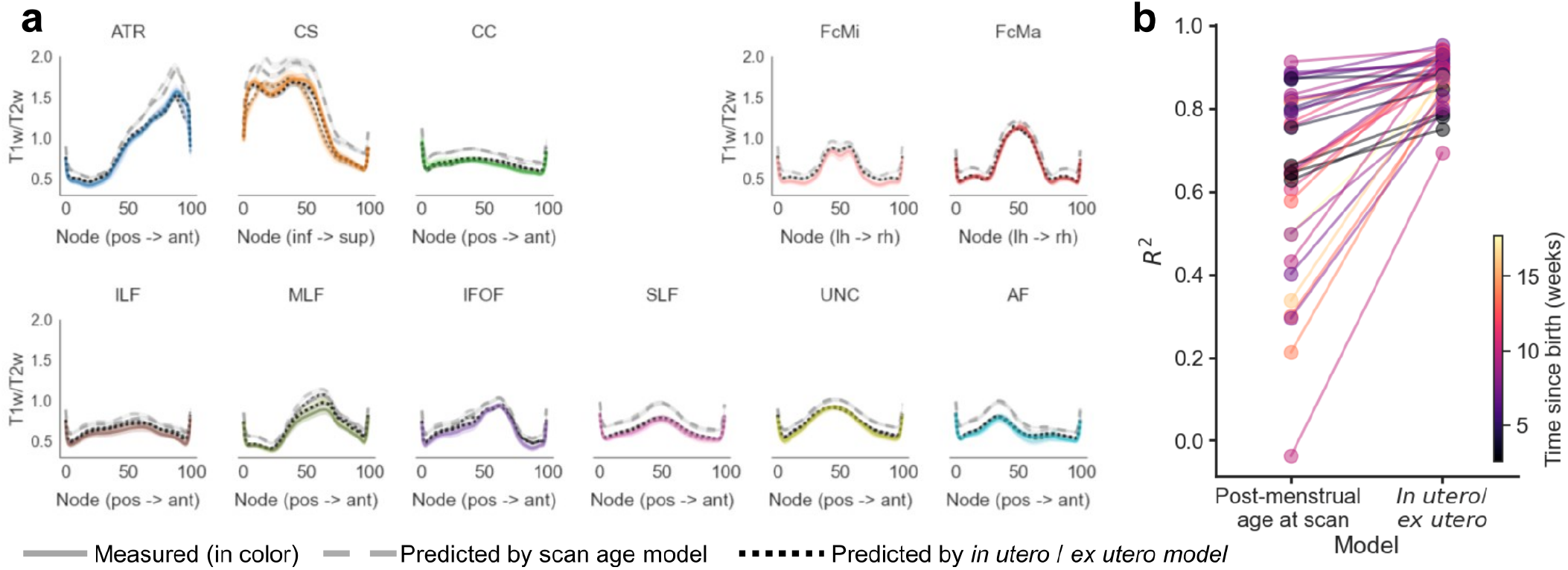
Model separating in utero and ex utero development best explains lower T1w/T2w at term-equivalent age in infants born preterm. a. Line graphs compare the measured T1w/T2w ratio in preterm infants at term-equivalent age (colored solid line) and the predicted T1w/T2w from two different sets of LMs fit on independent data. The first models used only post-menstrual age at scan as predictor (grey dashed line), whereas the second models accounted for time spend developing in utero and ex utero (black dotted line). Both hemispheres are presented in the same subplot, right hemisphere is shown in lighter hue. Shaded regions show 95% confidence intervals. b. Comparison of model fits between the post-menstrual age model and the in utero /ex utero model for T1w/T2w measured at all nodes across all bundles in the preterm infant’s second scan. Lines connect the same individual, indicating an overall increased model accuracy for the in utero /ex utero model. Colors indicate time spend developing ex utero for each individual. Abbreviations: ATR: anterior thalamic radiation, CS: cortico-spinal tract, CC: cingulum cingulate, FcMi: forceps minor, FcMa: forceps major, ILF: inferior longitudinal fasciculus, MLF: middle longitudinal fasciculus, IFOF: inferior frontal occipital fasciculus, SLF: superior longitudinal fasciculus, UNC: uncinate fasciculus, AF: arcuate fasciculus, pos: posterior, ant: anterior, inf: inferior, sup: superior, lh: left hemisphere, rh: right hemisphere

## Discussion

In the current study, we evaluated the impact that being born has on the development of T1w/T2w along bundles. To this end, we developed software for automated bundle identification in individual infant brains, and used it to analyse data from the dHCP (43). Using this large openly-available sample, we found that (i) T1w/T2w increases with increasing post-menstrual age at scan, (ii) there is slower T1w/T2w increase after birth (*ex utero*) than before birth (*in utero*), and (iii) slower *ex utero* T1w/T2w increase largely explains reduced T1w/T2w observed in infants born preterm. Taken together, these findings suggest that being born has fundamental impact on early-life development of the human white matter, which, in turn, can have negative consequences in cases where birth occurs too early during development.

Our finding of faster in utero than *ex utero* T1w/T2w increase is likely explained by the protective environment of the womb, with its consistent supply of nutrients, oxygen, warmth, and so forth. We speculate that the availability of nutrients may be a particularly important factor, as nutrition and myelination have been linked by animal models (10–12) and because human infants typically lose weight immediately after they are born (44), which illustrates the negative impact of being born on nutrition. Indeed, in humans, the link between nutrition and myelination is supported by work showing that the favorable nutritional composition of breast milk promotes myelination and cognitive development compared to formula (45–47). Further, recent work showed that even a delayed clamping of the umbilical cord, and the associated increased supply of iron, impacts myelination in infants (48). An interesting open question is the spatial specificity of the link between nutrition and myelination. Previous work on the benefits of breastfeeding on white matter development have provided conflicting results where either global (46) or regional specific (49) benefits were observed. In the current study, we find faster *in utero* T1w/T2w increase along the length of all included bundles, which aligns with the idea that nutrition and myelination are linked on a global scale. Another interesting open question that could be addressed in future work is how the beneficial effects of breastfeeding interact with the decelerating effect of birth, as exclusively breastfed babies typically lose more weight directly after birth, compared to their formula-fed peers (50). Further, infants typically regain their birth-weight by 2 weeks after birth (51), suggesting that nutritional deficiencies in newborns are relatively short-term. This raises the question of how long-lasting the decelerating impact of birth on white matter myelination is and whether myelination rates increase again once infants adapt to the new mode of feeding.

From the very moment infants are born, they are inundated with novel sensory input, particularly visual input. The theory of experience-dependent myelination, which was initially derived from animal models (7–9, 52), predicts that this increase in sensory signals leads to an immediate acceleration of myelination at birth, especially in bundles that convey sensory information. Here we find no evidence for this prediction, as T1w/T2w increase slows down at birth in all investigated bundles. This is surprising, as experience-dependent white matter changes are also found in humans (53). Interestingly, our data matches recent work on the myelination of visual cortex, which similarly showed that T1w/T2w in cortex is more strongly impacted by time spend developing *in utero* than *ex utero* (54). Our findings suggest that, to the degree that experience-dependent myelination starts occurring at birth, it is counteracted by other effects, such as nutritional changes. At later phase of development, experience-dependent myelination may become more dominant, explaining its key role in learning throughout the lifespan (55).

One limitation of the current study is the use of T1w/T2w to assess myelin development in infant white matter. To date, there is no histological work that directly validates the link between T1w/T2w and myelin content in the white matter early in life. Most prior studies examining the validity of using T1w/T2w to asses myelination used quantitative MRI measures (30–36) as a point of comparison. For these quantitative measures, such as quantitative R1, the link to myelin has been established (36), making them a meaningful benchmark. These studies find comparable test-retest reliability (56) and high consistency between T1w/T2w and quantitative measures in adult cortex (41), but limited consistency in adult white matter (57, 58). Indeed, the validity of using T1w/T2w in order to draw conclusions on adult white matter myelination has been further questioned by a recent histological study focused on the corpus callosum (59). However, importantly, the myelin content of infants’ white matter more closely aligns with adult cortex than adult white matter, as white matter myelination is only beginning at this age. Indeed, a recent study provided evidence supporting the usage of T1w/T2w to assess myelination in infant white matter (42) and our own data (Supplementary Fig. S4) shows high consistency between quantitative R1 and T1w/T2w across white matter bundles in a small sample of newborns. Nonetheless, more work is needed to validate the usage of T1w/T2w as a marker for myelin in different age groups and brain structures. This is particularly important, as most large scale data collection efforts, such as dHCP, include T1w and T2w but not quantitative measures, inducing a trade-off between large sample sizes and validated measures directly linked to myelin content. A second limitation of the current study is the predominant reliance on cross-sectional data. An alternative approach to assess the impact of being born on white matter myelination would have been to collect data from the same infants before and after birth. However, in order to fit accurate developmental slopes at least three measurements would have been needed prior to and after birth, which is currently infeasible, as in utero MRI is still in its infancy. Moreover, we successfully leveraged the models fit on cross-sectional data to predict T1w/T2w in a separate longitudinal sample, substantially increasing the confidence in our approach.

Our observations of faster *in utero* than *ex utero* development of T1w/T2w is particularly relevant in the context of preterm birth, as preterm infants spend more time developing *ex utero* compared to their full-term peers. Indeed, altered white matter structure is one of the most common negative consequences of preterm birth and has been linked to a multitude of developmental, neurological and cognitive disorders (60). The developmental delays that can result from preterm birth have also been shown to be long-lasting, persisting until school-entry (61) and beyond (62), whereas white matter abnormalities at term-equivalent age have been proposed as predictors for developmental outcomes in preterm infants (63). Post-mortem histology has suggested that the less mature preterm brain is more sensitive to its environment and more dependent on the protective environment of the womb for two reasons: i) Oligodendrocytes, the cells that generate myelin, are in a particularly fragile stage of their development and therefore susceptible to hypoxic injury, and ii) the infants’ vascular system is not fully developed leading to fluctuations in oxygen supply to the brain (13, 15, 16, 60, 64). Here we show that slower *ex utero* T1w/T2w growths largely explains the reduced T1w/T2w seen in preterm infants at term-equivalent age. Our results hence highlight general environmental factors that are independent of the infant’s maturity as a critical additional factor contributing to the white matter abnormalities seen in preterm infants. The present findings hence have implications for the care of preterm infants and predict that negative developmental outcomes may be reduced by closely matching the environment of infants born preterm to the environment they would have experienced in the womb. Indeed, larger nutrient intake after preterm birth has been shown to have a positive impact on white matter maturation at term-equivalent age, as well as on developmental outcomes (65), suggesting that compensating for birth-related reductions in nutrition can be beneficial for infants born preterm. Moreover, strikingly, it has even been shown that womb-like maternal sounds are beneficial for the development of the auditory cortex (66) and studies on the benefits of the maternal voice on white matter development are also underway (67).

In conclusion, the current study showed slower *ex utero* compared to *in utero* T1w/T1w growth in the first few weeks of life, suggesting that the sheltered environment of the womb is beneficial for white matter myelination. We also show that the reduced T1w/T2w seen in preterm infants at term-equivalent age is largely explained by the amount of time they spend developing *ex utero*. We hence hypothesize that closely matching the preterm infants environment to what they would have experienced in utero may promote white matter myelination and, in turn, developmental outcomes.

## Materials and Methods

### Data

We used data collected by the Developing Human Connectome project (dHCP) (43). Details regarding data acquisition parameters are provided directly by dHCP (http://www.developingconnectome.org/data-release/second-data-release/release-notes/). The dHCP also performed preprocessing for the anatomical (68) and diffusion-weighted data (69) and here we used the preprocessed data. We downloaded preprocessed data from the dHCP’s second data release from the study website (http://www.developingconnectome.org/). This release contained 490 sessions, from 445 individuals that included all of the data necessary for the current analyses (diffusion MRI + T1w + T2w). As a first quality assurance step, we removed all sessions that exceeded two standard deviations of the mean with regards to absolute motion and with regards to the amount of outlier slices replaced by FSL’s eddy tool during preprocessing. Five additional sessions were excluded during visual inspection due to obvious image artifacts or alignment errors. After tractography and bundle identification (see sections below), we conducted an additional quality assurance by excluding all sessions where one or more bundles contained 10 or less streamlines. The remaining data consisted of 402 sessions from 368 individuals. We divided this data into two separate samples. The “cross-sectional sample” contained data from 300 individuals scanned shortly after birth, including both infants born preterm (gestational age at birth less than 37 weeks) and full-term [135 females; gestational age at birth: range: 24.57 - 42.29 weeks, mean ± SD: 38.30 ± 3.50 weeks; post-menstrual age at scan: range: 29.29 - 44.71 weeks, mean ± SD: 39.90 ± 2.69 weeks, time between birth and scan: range: 0-16.71 weeks, mean ± SD: 1.60 ± 2.49 weeks]. The “longitudinal sample” contained 34 preterm infants scanned twice, once shortly after their preterm birth and again once they reached term-equivalent age [14 females; gestational age at birth: mean ± SD: 31.97 ± 2.96 weeks; post-menstrual age at first scan: mean ± SD: 34.34 ± 1.84 weeks; post-menstrual age at second scan: mean ± SD: 40.92 ± 1.34 weeks] as well as a group of full-term infants matched 1-on-1 to the preterm infants gender and post-menstrual age at second scan [14 females; gestational age at birth: mean ± SD: 40.08 ± 0.97 weeks; post-menstrual age at scan: mean ± SD: 40.92 ± 1.35 weeks].

### Tractography

DMRI tractography was performed in accordance with recent work (39), using MRtrix (70, 71). Voxel-wise fiber orientation distributions (FODs) were calculated with constrained spherical deconvolution (CSD). We used the Dhollander algorithm (72) to estimate three-tissue response functions, with the FA threshold set to 0.1 to account for the generally lower FA in the brains of newborns. We then computed FODs with multi-shell multi-tissue CSD (73) separately for the white matter and the CSF. The gray matter was not modeled separately, as white and gray matter do not have sufficiently distinct b-value dependencies to allow for a clean separation of the signals in newborns. Finally, we performed multi-tissue informed log-domain intensity normalization and generated a whole brain white matter connectome for each session. Tractography was optimized using the tissue segmentation from the anatomical MRI data (anatomically-constrained tractography, ACT (74)), which is particularly useful when gray and white matter cannot be separated in the FODs. For each connectome, we used probabilistic fiber tracking with the following parameters: algorithm: IFOD1, step size: 0.2 mm, minimum length: 4 mm, maximum length: 200 mm, FOD amplitude stopping criterion: 0.05, maximum angle: 15deg. Seeds for tractography were randomly placed within the gray/white matter interface (from anatomical tissue segmentation), which enabled us to ensure that streamlines reach the gray matter.

### Bundle identification with pyBabyAFQ

In order to evaluate white matter development along bundles in large samples, it is necessary to first identify the bundles in each individual infant’s native brain space in a systematic and automated way. Typically, automated tools for bundle identification in individuals have been developed for data from adults (75–78), and may not be suitable for the analysis of infant data. In our previous work, we developed baby automated fiber quantification in MATLAB (babyAFQ, (39)), an automated tool for bundle identification specifically catered to the infant brain. In order to analyse data at the scale of dHCP, we used a cloud computing environment. As scalable deployment benefits from fully open-source software, we ported babyAFQ to create a python implementation (pyBabyAFQ). Importantly, pyBabyAFQ is fully integrated into the pyAFQ software package (77), so that additional tools for plotting, bundle profile evaluation, and statistical analysis (e.g., AFQ-insight, https://github.com/richford/AFQ-Insight, (79)) can be applied after the bundles are identified. The newly developed pyBabyAFQ is fully open-source, simple to use, and straightforward to scale to very large datasets. The software identifies 20 white matter bundles in individual infant’s native brain space (9 in each hemisphere and 2 between-hemispheres, Supplementary Figure 2): the anterior thalamic radiation (ATR), cortico-spinal tract (CS), forceps major (FcMa), forceps minor (FcMi), arcuate fasciculus (AF), uncinate fasciculus (UNC), superior longitudinal fasciculus (SLF), cingulum cingulate (CC), inferior longitudinal fasciculus (ILF), inferior frontal occipital fasciculus (IFOF) and the middle longitudinal fasciculus (MLF). These bundles are identified using sets of anatomical ROIs that serve as way-points for each bundle. Critically, the way-point ROIs were defined in a newborn template (80), as previously described (39), and are transformed from this template space to each individual infant’s native brain space. A probabilistic atlas, which is also transformed from the newborn template space to the individual infant’s brain space, is used to determine which bundle is the better match for a given streamline in cases where a streamline crosses all way-points for multiple bundles. In each bundle, outlier streamlines were removed by finding streamlines whose trajectory deviated from the median trajectory of the bundle by more than 4 standard deviations. The Cloudknot software (81) was used to deploy both tractography (as described above) and bundle identification to the Amazon Web Services Batch service.

### Modeling T1w/T2w development

After identifying all bundles with pyBabyAFQ, we evaluated the development of T1w/T2w along the lengths of the bundles. First, we focused on the cross-sectional sample and fit linear models separately for each node in each bundle relating T1w/T2w either: (i) to post-menstrual age at scan (T1w/T2w ä 1 + post-menstrual age at scan), or (ii) to gestational age at birth, i.e., in utero development, and time between birth and scan, i.e., *ex utero* development (T1w/T2w ä 1 + gestational age at birth + time since birth). Models were compared using the Akaike information criterion (AIC). In the next step, we focused on the longitudinal sample, and compared T1w/T2w between: (i) preterm infants scanned right after birth, (ii) the same preterm infants scanned once they reached term-equivalent post-menstrual age, and (iii) a group of full-term infants matched to the preterm infants second scan. In our final analysis, we used the linear models fit on the cross-sectional sample to explain the T1w/T2w observed in the preterm infants from the longitudinal sample at their second scan. To compare the accuracy of the two linear models, we compared their model fits at all nodes in all bundles across individuals.

### Data and code availability

The data used in the current study is provided by the Developing Human Connectome project (dHCP, (43)) and can be downloaded through their website (http://www.developingconnectome.org/). The data were analyzed using open source software, including MRtrix (70, 71) and pyBabyAFQ, which we shared as a novel component of pyAFQ (https://yeatmanlab.github.io/pyAFQ (77)). Code that implemented the tractography pipeline, example code to perform bundle identification with pyBabyAFQ, and code used to generate the main figures of this manuscript are also made available in GitHub (https://github.com/EduNeuroLab/wmDevAroundBirth).

## ACKNOWLEDGMENTS

This research was funded through grant RF1MH121868 from the National Institute of Mental Health, through grant R01EY033835 from the National Eye Institute, and it was supported by the Deutsche Forschungsgemeinschaft (DFG, German Research Foundation) – project number 222641018 – SFB/TRR 135 TP C10 as well as by “The Adaptive Mind”, funded by the Excellence Program of the Hessian Ministry of Higher Education, Science, Research and Art. Credits from the Amazon Web Services research credits program supported cloud computing in this project. Data were provided by the developing Human Connectome Project, KCL-Imperial-Oxford Consortium funded by the European Research Council under the European Union Seventh Framework Programme (FP/2007-2013) / ERC Grant Agreement no. [319456]. We are grateful to the families who generously supported this trial. We thank Manjari Narayan for her help with data processing.

