## Supplementary Materials for "Human white matter myelination rate slows down at birth"

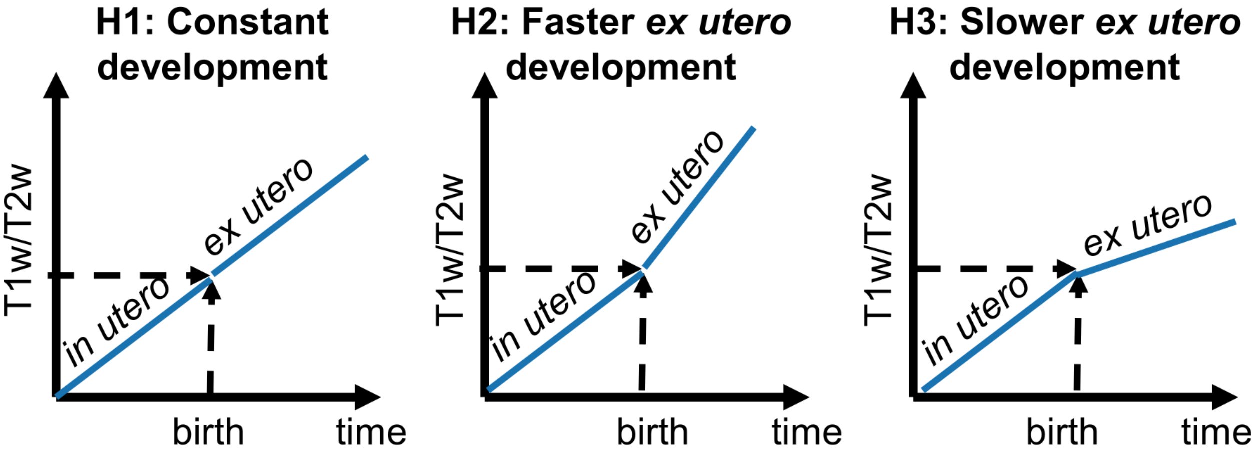


Supplementary Figure 1: Schematic of three hypotheses regarding relative rates of early life development of white matter T1w/T2w *in utero* and *ex utero*. H1: Development rate does not change at birth; H2: Development accelerates at birth; H3: Development decelerates at birth.

### pyBabyAFQ successfully identifies 20 white matter bundles in large samples

Evaluating early-life white matter development necessitates the identification of white matter bundles in individual infant's native brain space in a systematic and automated way. However, automated tools for bundle identification developed for data from adults [5, 6, 3, 1] may not be suitable for the analysis of

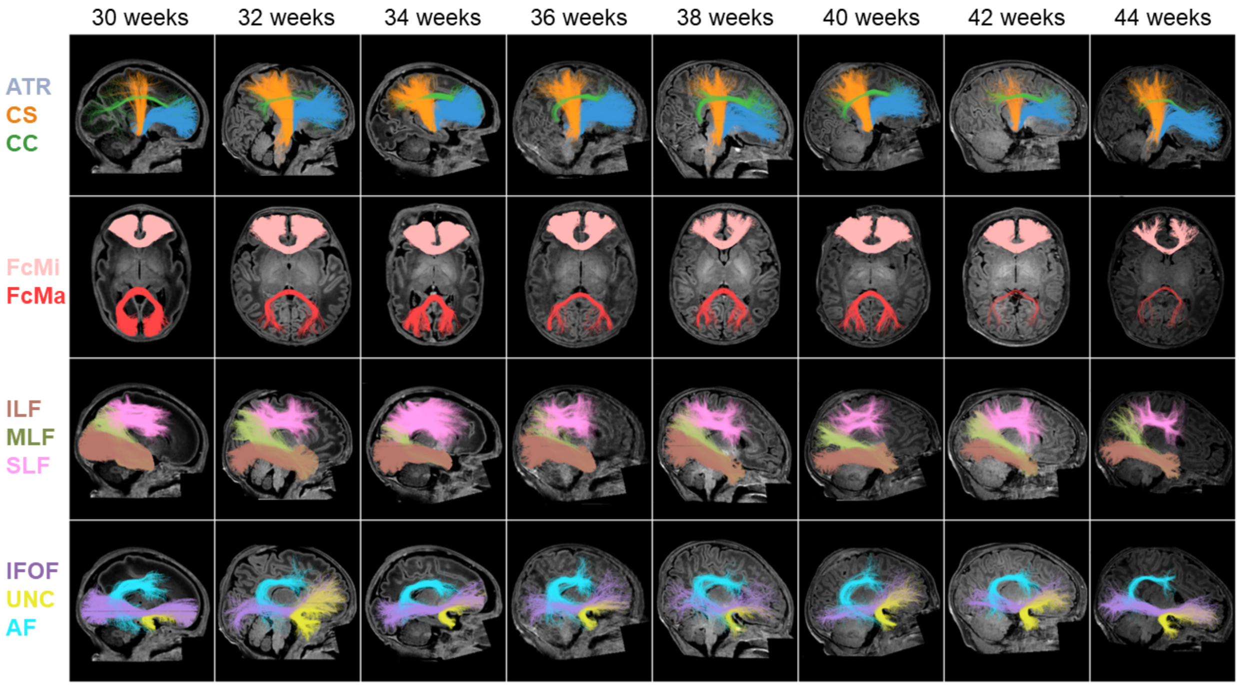


Supplementary Figure 2: Examples of bundles identified with pyBabyAFQ. Bundles are shown in eight randomly chosen individuals from dHCP scanned at post-menstrual ages ranging from 30 weeks (left) to 44 weeks (right). Abbreviations: ATR: anterior thalamic radiation, CS: cortico-spinal tract, CC: cingulum cingulate, FcMi: forceps minor, FcMa: forceps major, ILF: inferior longitudinal fasciculus, MLF: middle longitudinal fasciculus, IFOF: inferior frontal occipital fasciculus, SLF: superior longitudinal fasciculus, UNC: uncinate fasciculus, AF: arcuate fasciculus

infant data. As such, in our previous work, we developed baby automated fiber quantification (babyAFQ, [2]) an automated tool for bundle identification specifically catered to the infant brain. To analyze dHCP data in a cloud computing environment, we ported the software and create a python implementation of babyAFQ (pyBabyAFQ), which we integrated into the pyAFQ software package [3]. The newly developed pyBabyAFQ is fully open-source, simple to use, and straightforward to scale to very large datasets (see also [4]). The software identifies 20 white matter bundles in individual infant's native brain space (9 in each hemisphere and 2 between-hemispheres, Supplementary Figure 2). In order to validate pyBabyAFQ, we tested it on a small sample of newborns (N=9) collected at Stanford University for which manually identified 'gold-standard' bundles are available (these data were previously published in [2]; see therein for details on the sample, the acquisition, and manual bundle delineation). We used the dice coefficient to quantify the spatial overlap between these manual bundles and bundles identified automatically using three different tools: (i) classical AFQ developed for adult data in MATLAB [5], (ii) babyAFQ in MATLAB (as described in [2]), and (iii) pyBabyAFQ. We found that the MATLAB and python implementations of babyAFQ perform similarly and show more spatial overlap with manual bundles than classical AFQ (repeated measures analysis of variance (rmANOVA) with AFQ version and bundles as factors: main effect of AFQ-version: $F(2,16)=448.21$, $p<0.0001$; post-hoc comparisons: AFQ and babyAFQ: $p<0.0001$, AFQ and pyBabyAFQ: $p<0.0001$, babyAFQ vs pyBabyAFQ: $p=0.77$; Supplementary Figure 3). Next, we applied pyBabyAFQ to data from the dHCP project and found that it successfully identifies all 20 bundles in 95% of all included data sets, suggesting that pyBabyAFQ is reliably scalable for the analysis of large samples.

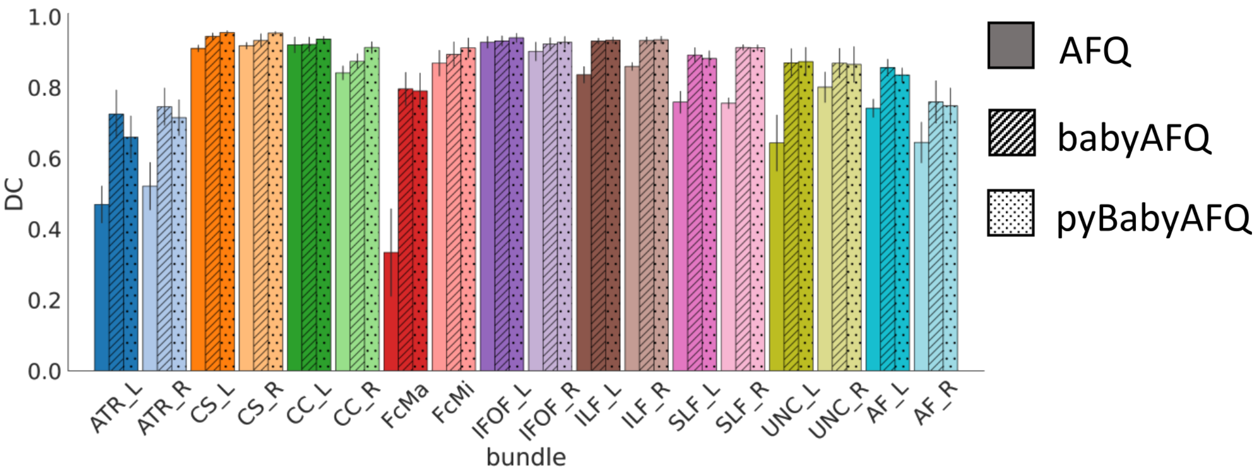


Supplementary Figure 3: PyBabyAFQ is well-suited for bundle identification in infants. Comparison of AFQ and babyAFQ performances in identifying each bundle in newborns (n=9) relative to manually defined (gold-standard) bundles. Overlap between automatically and manually defined bundles is evaluated using the dice coefficient (DC), which reveals higher performance for both the MATLAB and the python implementation of babyAFQ than AFQ. Bars show mean dice coefficient ± standard error across participants. Abbreviations: ATR: anterior thalamic radiation, CS: cortico-spinal tract, CC: cingulum cingulate, FcMi: forceps minor, FcMa: forceps major, ILF: inferior longitudinal fasciculus, MLF: middle longitudinal fasciculus, IFOF: inferior frontal occipital fasciculus, SLF: superior longitudinal fasciculus, UNC: uncinate fasciculus, AF: arcuate fasciculus, L: left hemisphere, R: right hemisphere

### T1w/T2w correlates with quantitative R1

We also used the Stanford dataset described above to validate the usage of T1w/T2w as a proxy for myelination in the infant white matter. To this end, we compared T1w/T2w to quantitative R1 measures available in that sample (see [2]). We found that T1w/T2w strongly correlates with R1 across white matter bundles in neonates (Pearson $r^2 = 0.94$, Supplementary Figure 4). Lower correlations with R1 were found for commonly used diffusion metrics such as MD ($r^2 = 0.86$, Supplemental Figure 4), and FA ($r^2 = 0.08$, Supplementary Figure 4). The high correlation with R1 supports the feasibility of using T1w/T2w as a myelin-sensitive imaging contrast when R1 data is not available.

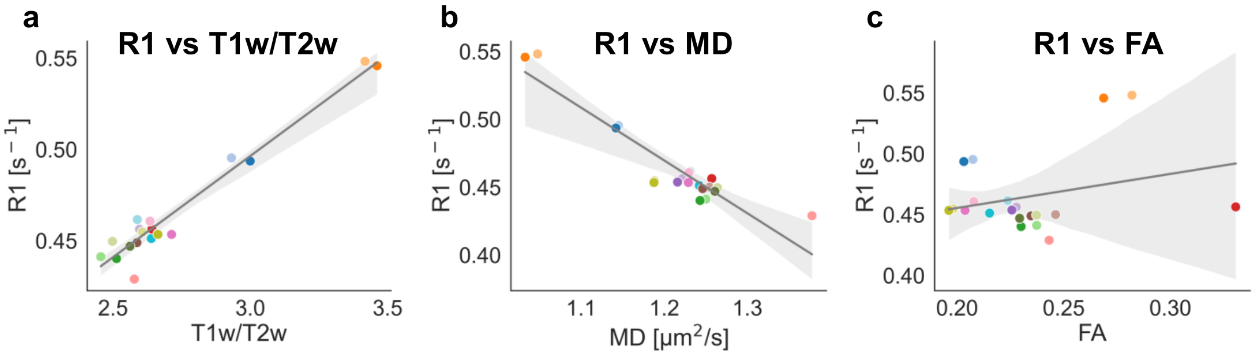


Supplementary Figure 4: Myelin-weighted quantitative MRI measurements of the R1 time-constant in 9 newborns (age 8–37 days) are compared to T1w/T2w (a) and to dMRI-derived mean diffusivity (MD) (b), and fractional anisotropy (FA) (c) across 20 white matter bundles. The correlation of R1 is remarkably high with T1w/T2w ($r^2 = 0.94$), is lower with MD ($r^2 = 0.86$), and even lower with FA ($r^2 = 0.08$). The high correlation with R1 strongly supports the feasibility of using T1w/T2w as a myelin-sensitive imaging contrast. Abbreviations: MD: mean diffusivity, FA: fractional anisotropy

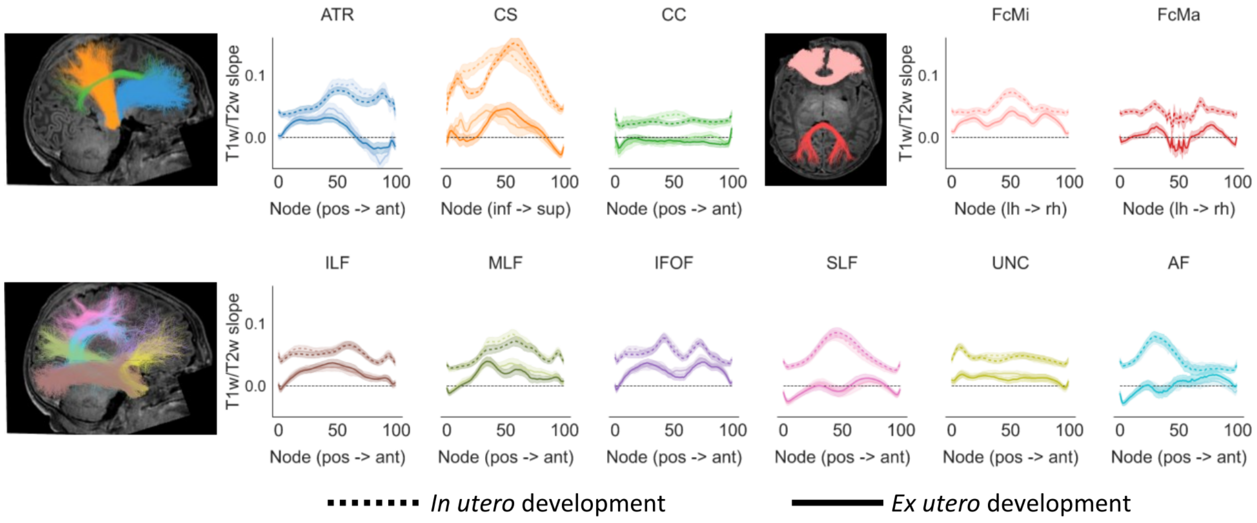


Supplementary Figure 5: The observation of faster *in utero* than *ex utero* T1w/T2w increase remains when preterm born infants are excluded. Here, we reproduced the plots from Figure 2 while only including subjects born at gestational age of 37 weeks or later from the cross-sectional samples (n=234). The conclusions reached in Figure 2 in the main text are also supported by this sub-set of the data. Abbreviations: ATR: anterior thalamic radiation, CS: cortico-spinal tract, CC: cingulum cingulate, FcMi: forceps minor, FcMa: forcepts major, ILF: inferior longitudinal fasciculus, MLF: middle longitudinal fasciculus, IFOF: inferior frontal occipital fasciculus, SLF: superior longitudinal fasciculus, UNC: uncinate fasciculus, AF: arcuate fasciculus, pos: posterior, ant: anterior, inf: inferior, sup: superior, lh: left hemisphere, rh: right hemisphere

| | scan age model | | *in utero* / *ex utero* model | | |
|---|---|---|---|---|---|
| bundle | $R^2$ | T1w/T2w slope | $R^2$ | T1w/T2w slope *in utero* | T1w/T2w slope *ex utero* |
| AF_LH | 0.31 | 0.03 | 0.38 | 0.03 | 0.02 |
| AF_RH | 0.33 | 0.03 | 0.40 | 0.03 | 0.02 |
| ATR_LH | 0.43 | 0.04 | 0.51 | 0.04 | 0.02 |
| ATR_RH | 0.35 | 0.04 | 0.42 | 0.04 | 0.02 |
| CC_LH | 0.30 | 0.02 | 0.40 | 0.02 | 0.01 |
| CC_RH | 0.23 | 0.02 | 0.32 | 0.03 | 0.01 |
| CS_LH | 0.59 | 0.08 | 0.68 | 0.09 | 0.05 |
| CS_RH | 0.52 | 0.08 | 0.60 | 0.08 | 0.05 |
| FcMi | 0.41 | 0.03 | 0.45 | 0.03 | 0.02 |
| FcMa | 0.36 | 0.03 | 0.42 | 0.03 | 0.02 |
| IFOF_LH | 0.51 | 0.04 | 0.55 | 0.04 | 0.03 |
| IFO_RH | 0.52 | 0.04 | 0.55 | 0.04 | 0.03 |
| ILF_LH | 0.50 | 0.04 | 0.53 | 0.04 | 0.03 |
| ILF_RH | 0.51 | 0.04 | 0.55 | 0.04 | 0.03 |
| MLF_LH | 0.46 | 0.04 | 0.50 | 0.04 | 0.03 |
| MLF_RH | 0.46 | 0.04 | 0.50 | 0.04 | 0.03 |
| SLF_LH | 0.43 | 0.04 | 0.53 | 0.04 | 0.02 |
| SLF_RH | 0.40 | 0.04 | 0.50 | 0.04 | 0.02 |
| UNC_LH | 0.53 | 0.04 | 0.59 | 0.04 | 0.02 |
| UNC_RH | 0.55 | 0.04 | 0.61 | 0.04 | 0.03 |

Supplementary Table 1: Details on the model coefficients (T1w/T2w slopes) and model fits for each bundle, separately for the post-menstrual age at scan model and the *in utero* / *ex utero* model. Values are the mean across all nodes in a bundle. Abbreviations: ATR: anterior thalamic radiation, CS: cortico-spinal tract, CC: cingulum cingulate, FcMi: forceps minor, FcMa: forceps major, ILF: inferior longitudinal fasciculus, MLF: middle longitudinal fasciculus, IFOF: inferior frontal occipital fasciculus, SLF: superior longitudinal fasciculus, UNC: uncinate fasciculus, AF: arcuate fasciculus, LH: left hemisphere, RH: right hemisphere
